## Supplementary Figures for "The structural context of PTMs at a proteome wide scale"

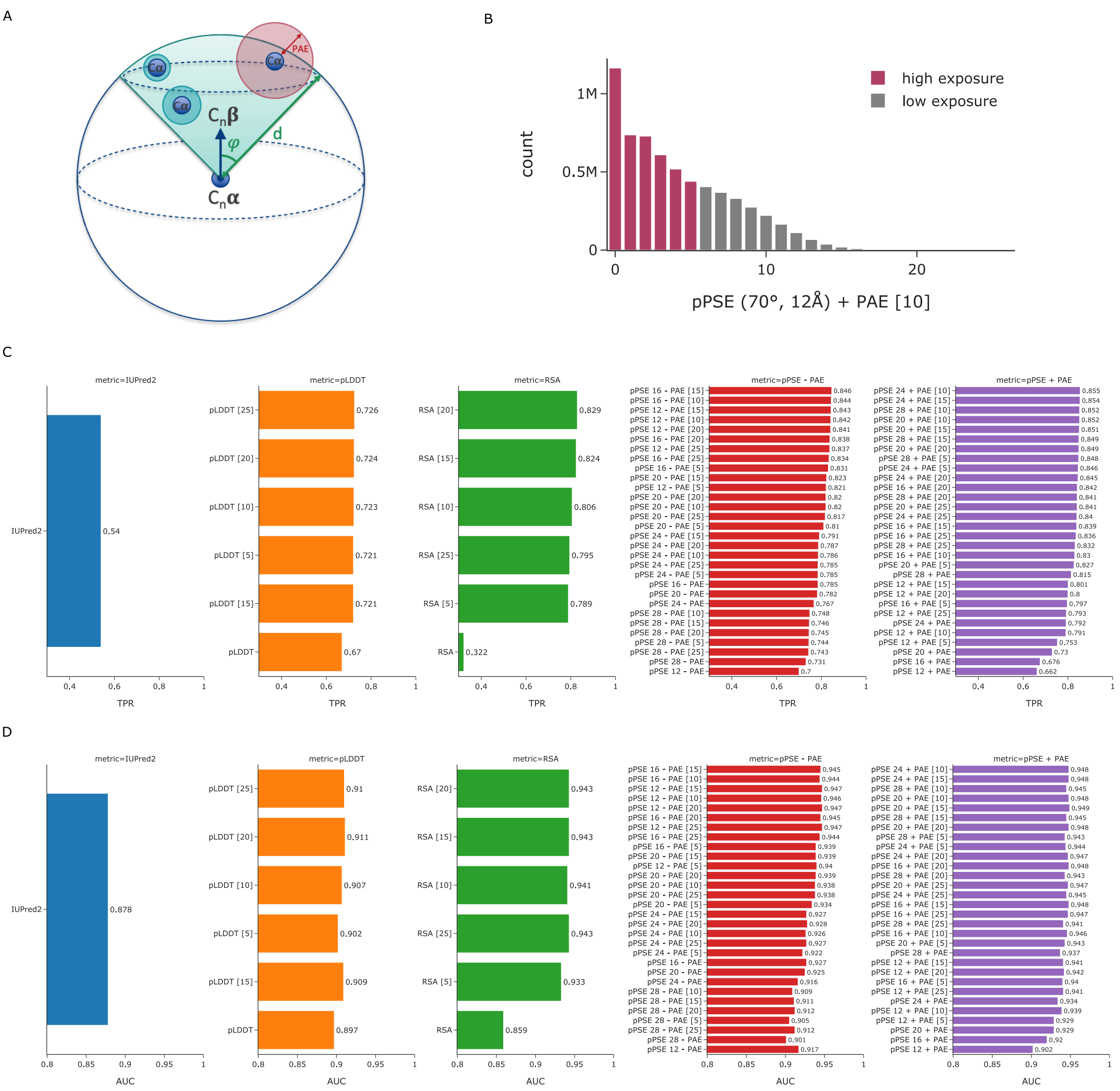


**Supplementary Figure 1. Estimation of amino acid side chain exposure and intrinsically disordered regions (IDRs). (A)** Visualization of the strategy to calculate the prediction-aware part-sphere exposure (pPSE). **(B)** Distribution of pPSE values across all amino acids in structured protein regions (non-IDRs). **(C)** Parameter screen to evaluate the ability of different metrics to predict IDRs based on the true positive rate (TPR) at a 5% false positive rate (FPR). **(D)** Parameter screen to evaluate the ability of different metrics to predict IDRs based on the area under the curve (AUC). The numbers in square brackets behind each metric indicate the smoothing windows that were used.

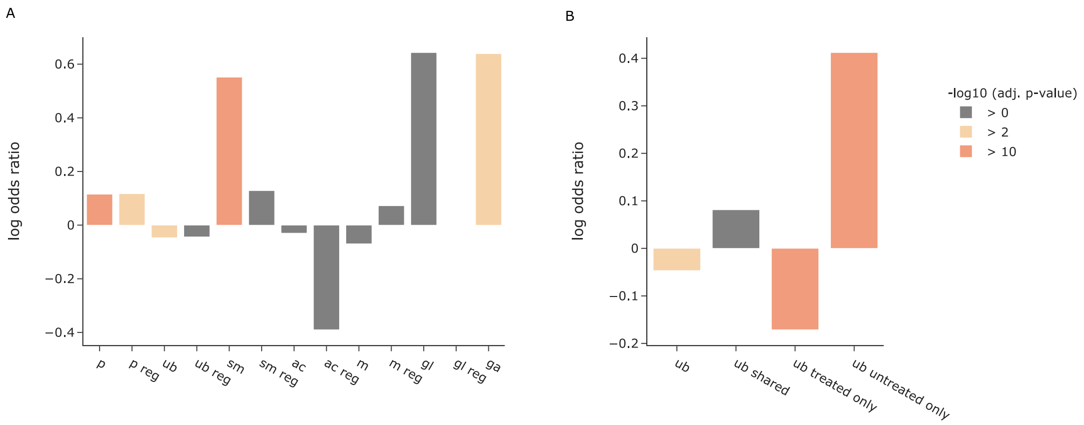


**Supplementary Figure 2. Enrichment analysis of PTMs in amino acids with high versus low side chain exposure. (A)** Enrichment of different PTMs annotated in the PhosphoSitePlus database in amino acids with side chains of high side chain exposure within structured regions. PTMs are abbreviated as follows: phosphorylations (p), ubiquitinations (ub), sumoylations (sm), acetylations (ac), methylations (m) and the glycosylations O-GalNAc (gl) and O-GlcNAc (ga). **(B)** Enrichment of ubiquitinated lysines annotated in PhosphoSitePlus versus ubiquitinations detected in a dataset treated with proteasome inhibitor or untreated.

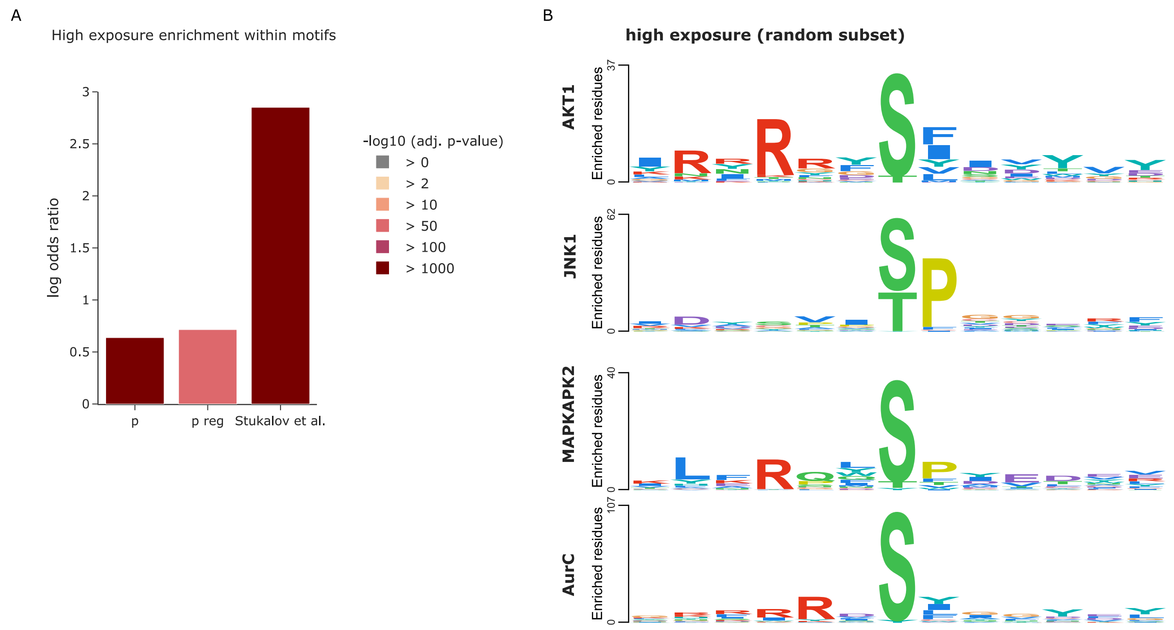


**Supplementary Figure 3. Exploiting the 3D context of kinase phosphorylation motifs. (A)** Enrichment of phosphorylations in kinase motifs with amino acids of high side chain exposure compared to all possible kinase motif occurrences in structured regions. **(B)** Sequence logos for different kinases based on a random subset of high-exposure sites, comprising the same number of sites as compared to the low exposure set. The PSSMSearch tool (Krystkowiak et al., 2018) was used with a log odds scoring method (O'Shea et al., 2013).

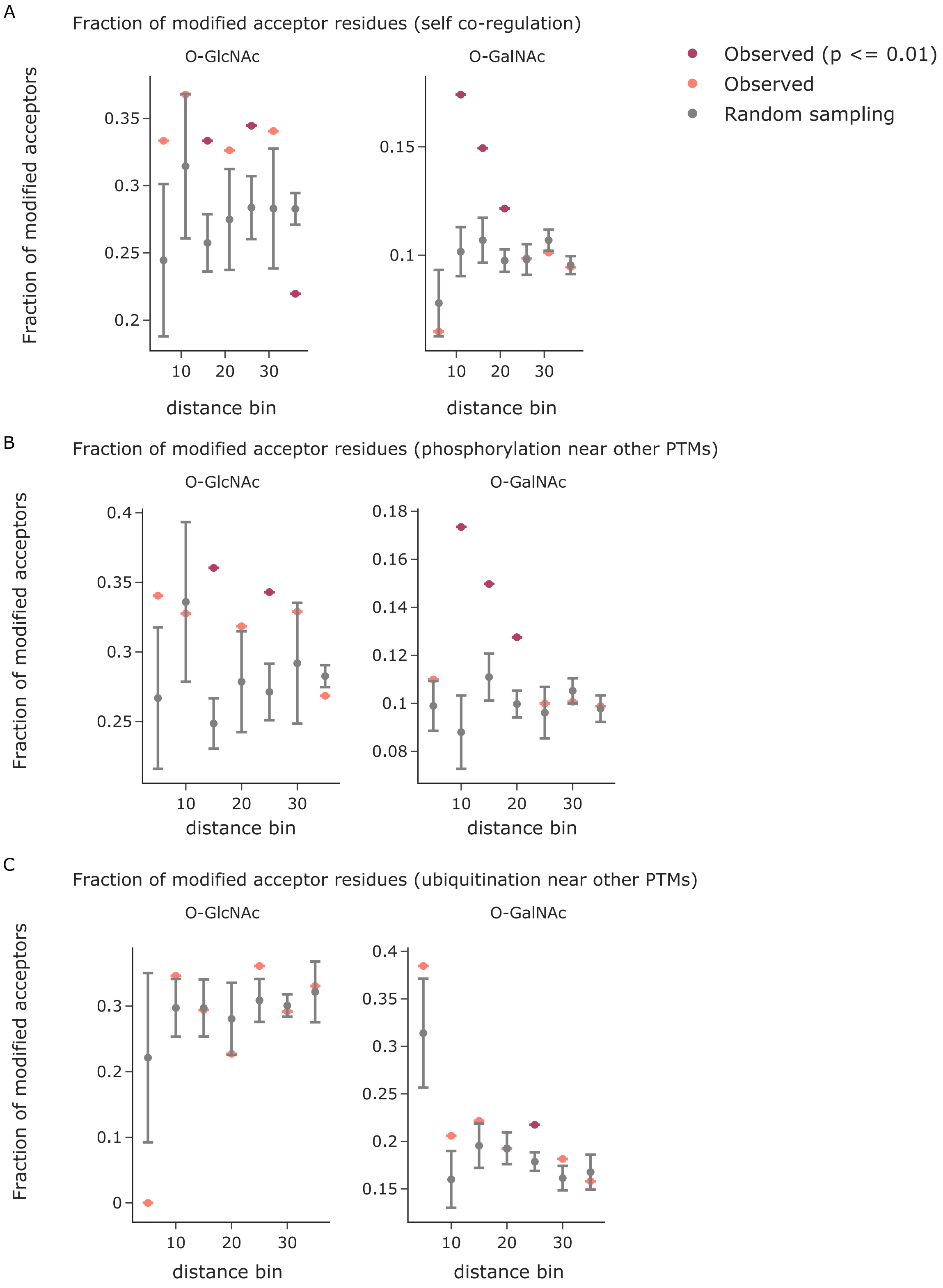


**Supplementary Figure 4. PTM proximity analysis in 3D. (A)** The fraction of modified PTM acceptor residues is shown as a function of the 3D distance to a given modified amino acid in Å. Observed values (indicated in red when statistically significant and colored in salmon otherwise) are compared to the mean of five random samples including the same number of modified PTM sites (grey). Error bars indicate one standard deviation. The x-axes are divided in distance bins ranging from each previous bin to the indicated cutoff in Å. **(B)** The fraction of modified phospho-acceptor residues is shown as function of the 3D distance to a given modified amino acid in Å. **(C)** The fraction of ubiquitinated lysines is shown as function of the 3D distance to a given modified amino acid in Å. The smallest bin shows competition for the same central lysine residue.
